# CDK6-mediated endothelial cell cycle acceleration drives arteriovenous malformations in hereditary hemorrhagic telangiectasia

**DOI:** 10.1101/2023.09.15.554413

**Authors:** Sajeth Dinakaran, Haitian Zhao, Yuefeng Tang, Zhimin Wang, Santiago Ruiz, Aya Nomura-Kitabayashi, Lionel Blanc, Marie E. Faughnan, Philippe Marambaud

## Abstract

Increased endothelial cell (EC) proliferation is a hallmark of arteriovenous malformations (AVMs) in hereditary hemorrhagic telangiectasia (HHT). The underlying mechanism and disease relevance of this abnormal cell proliferative state of the ECs remain unknown. Here, we report the identification of a CDK6-driven mechanism of cell cycle progression deregulation directly involved in EC proliferation and HHT vascular pathology. Specifically, HHT mouse liver ECs exhibited defects in their cell cycle control characterized by a G1/S checkpoint bypass and acceleration of cell cycle speed. Phosphorylated retinoblastoma (p-RB1)—a marker of G1/S transition through the restriction point—significantly accumulated in ECs of HHT mouse retinal AVMs and HHT patient skin telangiectasias. Mechanistically, ALK1 loss of function increased the expression of key restriction point mediators, and treatment with palbociclib or ribociclib, two CDK4/6 inhibitors, blocked p-RB1 increase and retinal AVMs in HHT mice. Palbociclib also improved vascular pathology in the brain and slowed down endothelial cell cycle speed and EC proliferation. Specific deletion of *Cdk6* in ECs was sufficient to protect HHT mice from AVM pathology. Thus, CDK6-mediated endothelial cell cycle acceleration controls EC proliferation in AVMs and is a central determinant of HHT pathogenesis. We propose that clinically approved CDK4/6 inhibitors have repurposing potential in HHT.

## INTRODUCTION

Hereditary hemorrhagic telangiectasia (HHT) is an autosomal dominant vascular dysplasia affecting 1 in 5000 individuals and characterized by focal development of arteriovenous malformations (AVMs) in several tissues ^1^. Affected organs and tissues include the liver, lungs, and brain—with up to 20% of HHT patients exhibiting cerebral vascular pathology ^2^—as well as the mucosa and skin, where AVMs lead to small, enlarged, and superficial clustered vessels called telangiectasias. AVMs originate from dilations of defective blood capillaries that allow the formation of high flow shuts between an artery and a vein. Telangiectasias in the oronasal or intestinal mucosa lead to epistaxis and intestinal bleeding, respectively, while AVMs in the brain, liver, and lungs increase the risk of stroke and internal hemorrhage, and consecutive anemia, organ failure, and cardiac complications ^3–5^.

Loss of function (LOF) heterozygous germline mutations in HHT families are found in different genes that all share the same TGF-β signaling pathway ^6,7^. These genes include *ENG* (coding for endoglin/ENG and leading to HHT type 1, HHT1), *ACVRL1* (activin receptor-like kinase 1, ALK1; HHT2), and *SMAD4* (mothers against decapentaplegic homolog 4, Smad4; juvenile polyposis (JP)-HHT) ^8–10^. ALK1 is a BMP type I receptor that forms cell surface protein complexes with a BMP type II receptor and the accessory receptor ENG to bind BMP9 and BMP10 circulating ligands ^11–13^. ALK1-ENG receptor activation induces phosphorylation of the signal transducers, Smad1 and 5, to lead to the formation of Smad1/5-Smad4 complexes that translocate into the nucleus to control specific gene expression programs involved in vascular development and maintenance ^14^.

Inheritance of an HHT germline mutation does not necessarily lead to vascular pathology in patients, and when lesions occur, they manifest focally and generally progress over time ^14^. A model has emerged suggesting that both bi-allelic LOF of the HHT genes and angiogenic stimulation are required for disease manifestation. Key findings in HHT patients include the identification of somatic mutations in AVMs that lead to bi-allelic LOF in *ACVRL1* or *ENG* ^15^, and the observation that the anti-VEGFA antibody bevacizumab has possible therapeutic value in HHT ^16,17^. Early work in mice and zebrafish deficient for ENG, ALK1, Smad4, or BMP9/10 confirmed this model by showing that only full inactivation of ALK1 signaling during postnatal angiogenic development, or in adult mice challenged with healing, inflammatory, or VEGFA overexpression conditions, could reliably develop vascular pathologies ^13^. In this context, the mouse postnatal retinal angiogenesis model gained significant momentum in HHT AVM research. The main advantages of this model include a well-defined anatomical and temporal formation of the AVMs, the short time frame for disease development (about one week), and the high disease penetrance: nearly 100% when pups are treated with anti-BMP9 and anti-BMP10 blocking antibodies, hereafter referred to as the BMP9/10 immunoblocking (BMP9/10ib) model^18–20^.

HHT pathogenesis originates in the ECs. ALK1 and ENG are highly expressed in the endothelium, and conditional deletion of *Acvrl1*, *Eng*, *Smad4*, or *Smad1/5* in ECs is sufficient to cause AVMs ^13^. Defects in this pathway then lead to a complex array of signaling and cell behavior changes in the AVMs, not only in ECs but also in mural cells, such as smooth muscle cells (SMCs) that muscularize the AVMs ^7^. These changes include increased EC proliferation and size, and defects in EC migration against the blood flow ^21,22^. Aberrant EC proliferation is a particularly interesting aspect of the pathogenic process because it could be required for the structural development and maintenance of the AVMs. Whether EC proliferation is a cause or a consequence of AVM development and whether its inhibition could block AVM pathology remain unknown.

In adult vessels without angiogenic stimulation, most ECs are in a non-cycling, G0 quiescent state, characterized by the absence of proliferation ^14,23,24^. Reactivating the cell cycle in quiescent ECs requires G0 and G1 phase exit and successful transition through the restriction (R) point before entering the S phase. Cell cycle progression requires the inactivation of the retinoblastoma protein (RB1), an inhibitor of the transcription factors E2Fs. During the R point, RB1 inactivation is governed by its sequential phosphorylation by CDK4 or CDK6 and CDK2 ^25,26^. BMP9/10-ALK1-ENG signaling plays an important role in EC quiescence, and its disruption was proposed to contribute to the disease process ^14^. Evidence suggests that EC proliferation in HHT might be associated with cell cycle deregulations. Indeed, ALK1 and ENG activity controls endothelial cell cycle arrest *in vitro* ^27,28^ and *in vivo* ^29^, and fluid shear stress limits EC proliferation *via* ALK1-ENG signaling and cell cycle inhibition ^30–33^, a process that might be deregulated in the AVMs. Furthermore, we and others have reported that the overactivation of mTOR, an essential nutrient sensor that coordinates cell growth and cell cycle ^34^, is observed in ECs of the AVMs in different HHT models ^20,29^ and HHT2 patients ^35^. However, the exact mechanism driving EC loss of quiescence, abnormal cell cycle progression, and EC proliferation in HHT remains elusive. Here, we asked how the cell cycle is deregulated in ECs of HHT mice and patient skin biopsies and whether this deregulation is a key determinant in EC proliferation and AVM pathogenesis.

## RESULTS

### Endothelial G1/S checkpoint bypass and cell cycle acceleration accompany EC hyperproliferation in HHT mice

BMP9 and BMP10 promote EC quiescence ^36,37^, and retinal ECs proliferate in *Eng*, *Alk1*, *Smad4, and Bmp10* deficient mice ^38–43^. Similarly, in postnatal day 6 (P6) retinas of the BMP9/10ib model (**Fig. 1A**), we found an increased number of arterial and venous ECs (ERG^+^ cells) in the vasculature [isolectin B4 (IB4)-stained] of the AVM-containing peri-optic nerve area and in the mid-plexus area, compared to control mouse retinas (**Figs. 1B-D**). EdU staining in P6 retinas showed an increase in ERG^+^ EdU^+^ cells in both the AVM-containing peri-optic nerve area and hypervascularized mid-plexus of the BMP9/10ib mice, compared to control mice (**Figs. 1B, 1E, 1F**), confirming that ECs undergo hyperproliferation. EdU-stained liver ECs were then isolated to determine their cell cycle phase distribution by flow cytometry. Results showed that while the number of cells in G1 was decreased, the number of cells in the S phase was increased in BMP9/10ib mice, compared to controls (**Figs. 2A, 2B**). Mean fluorescence intensity (MFI) of S phase cells, which indicates the intra-S phase rate of DNA synthesis and which is inversely related to S phase duration ^44^, was measured and was found to be significantly increased in BMP9/10ib mouse ECs, compared to control ECs (**Figs. 2C, 2D**), suggesting that cycling speed is increased in BMP9/10ib mice. To confirm the mechanism leading to EC proliferation, we turned to the inducible two-color reporter histone H2B-fluorescent timer (iH2B-FT) mice that allow resolving cell cycle speed in a single snapshot measurement ^45^. When newly synthesized, the color-changing FT protein emits blue fluorescence and then red fluorescence during protein maturation. This unique fluorescent model allows the identification and quantification of faster cycling cells. iH2B-FT;BMP9/10ib mice were generated, and liver ECs were isolated at P8 for live-cell analysis by flow cytometry for blue and red fluorescence analysis. ECs from iH2B-FT;BMP9/10ib livers showed a robust, population-wide shift toward the blue fluorescence axis (**Fig. 2F**) and contained a significantly higher percentage of fast-cycling ECs (emitting blue fluorescence) compared to iH2B-FT;PBS control littermates (**Fig. 2G**), demonstrating that endothelial cell cycle speed is significantly increased in BMP9/10ib mouse ECs. Together, these results show that increased EC proliferation was accompanied by cell cycle defects characterized by a G1/S checkpoint bypass and cell cycle speed acceleration in ECs.

**Fig. 1.**
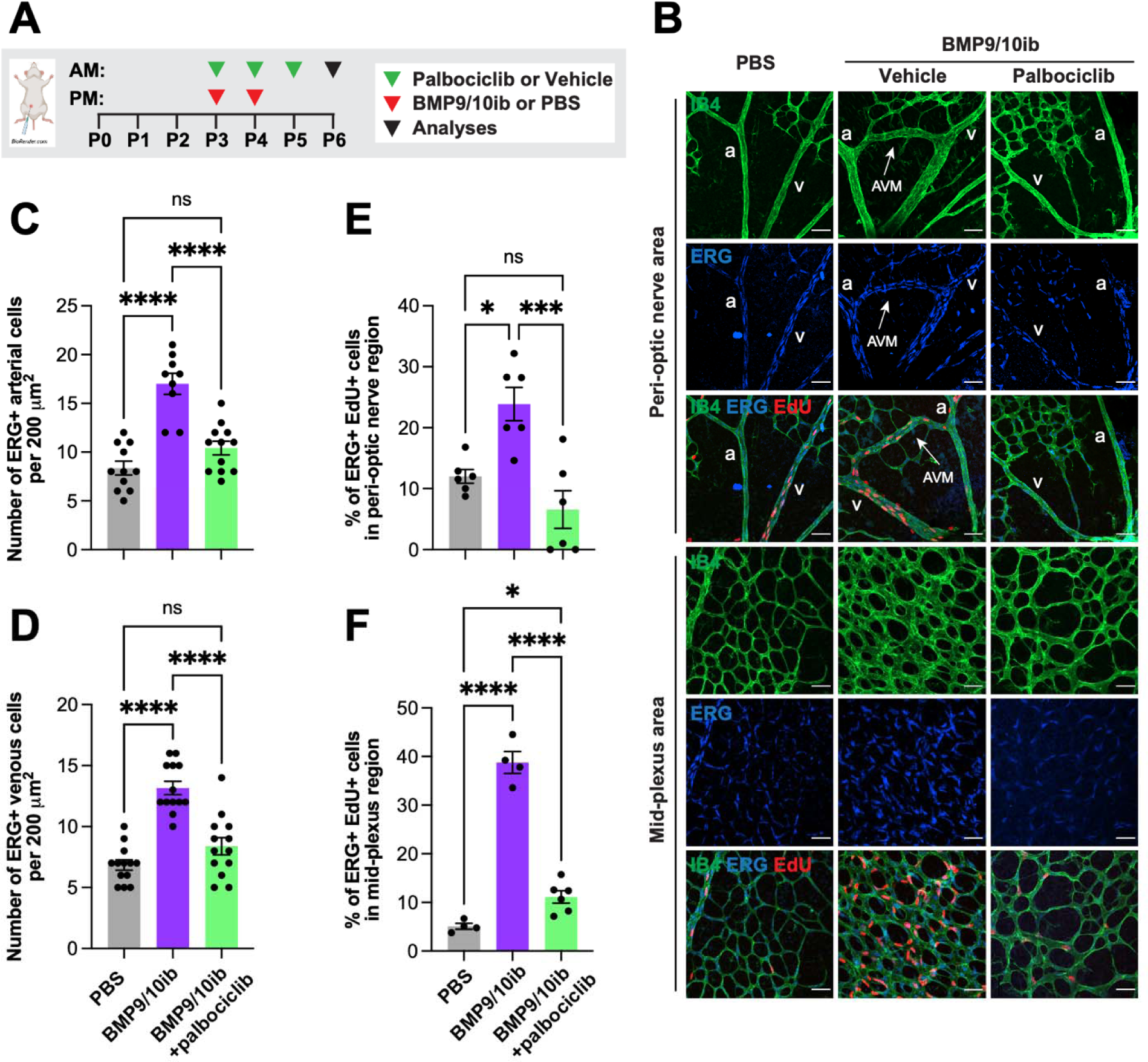
EC number and proliferation in HHT mice: Effect of palbociclib. (**A**) Schematic representation of the i.p. injection protocol. Arrowheads indicate the postnatal days (P) of injection (AM or PM). Pups were euthanized at P6 for analysis. (**B**) Representative staining using isolectin B4 (IB4, green), and of ERG (blue) and EdU (red) in the peri-optic nerve and mid-plexus regions of retinas from PBS controls, vehicle-treated BMP9/10ib mice, and palbociclib-treated BMP9/10ib mice. Arrows denote AVMs; a, artery; v, vein. Scale bars, 50 μm. (**C and D**) Scatter plots showing the total number of ERG^+^ cells per 200 μm^2^ in arteries (**C**) and veins (**D**) in the peri-optic nerve region across three groups: PBS (n=11), vehicle-treated BMP9/10ib (n=9), and palbociclib-treated BMP9/10ib (n=12) mice. (**E and F**) Scatter plots showing ERG^+^ EdU^+^ cells in the peri-optic nerve (**E**) and mid-plexus (**F**) regions across three groups: PBS (n=4-6), vehicle-treated BMP9/10ib (n=4-6), and palbociclib-treated BMP9/10ib (n=6) mice. Data represent individual retinas and mean ± SEM, one-way ANOVA with Tukey’s multiple comparisons test. ns, not significant; *P < 0.05, ***P < 0.001, ****P < 0.0001.

**Fig. 2.**
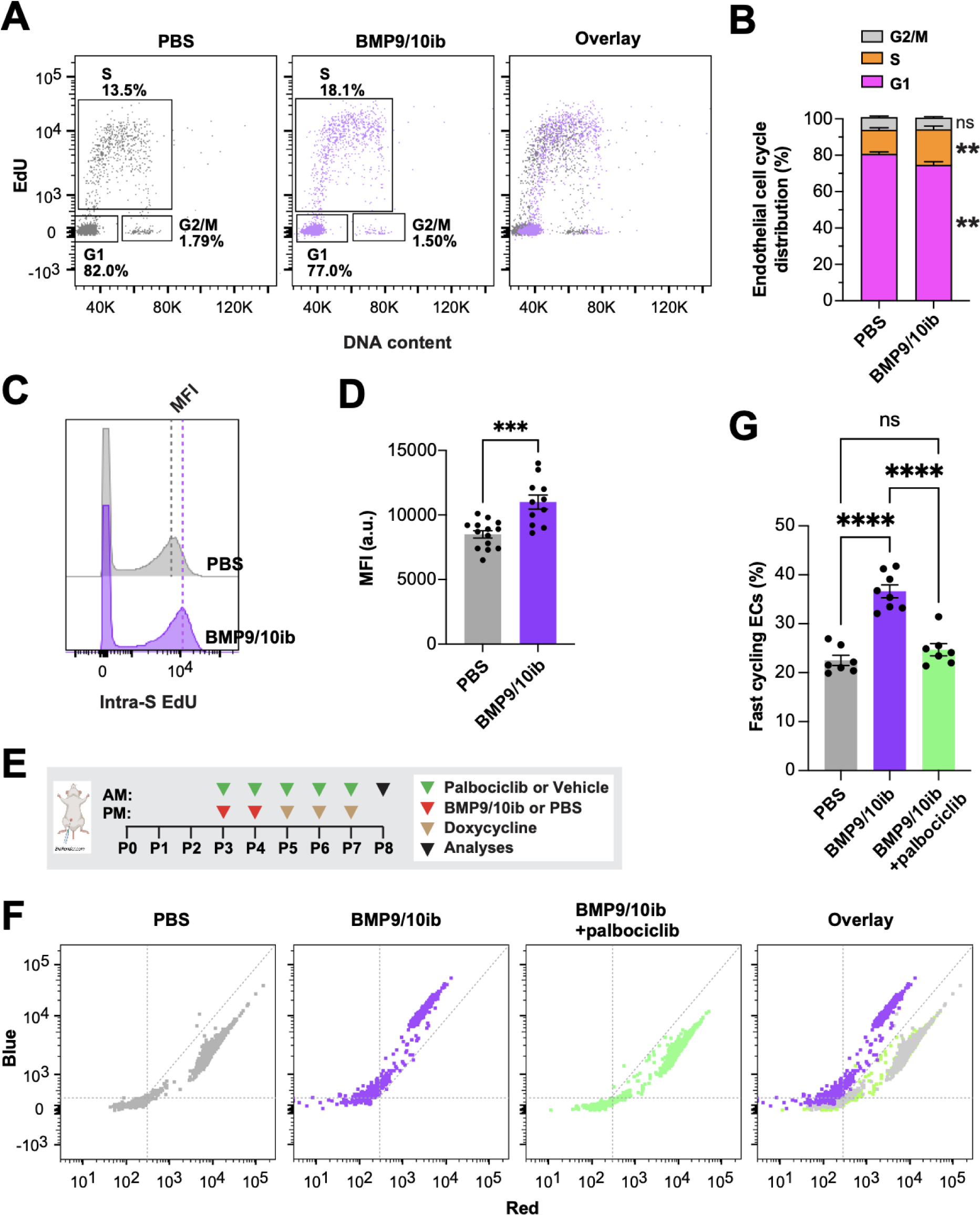
Endothelial cell cycle progression and speed in HHT mice: Effect of palbociclib. (**A and B**) Representative flow cytometry plots (**A**) and quantification (**B**) of cell cycle phase distribution of ECs isolated from livers of P6 and P8 PBS (n=14) and BMP9/10ib (n=11) mice. (**C and D**) Representative flow cytometry (**C**) and quantification (**D**) of cells analyzed as in (**A**) showing S phase duration measured as intra-S phase of EdU incorporation [intra-S EdU mean fluorescence intensity (MFI)]. Data in (**B**) and (**D**) represent mean ± SEM, unpaired t-test. (**E**) Schematic representation of the i.p. injection protocol. Arrowheads indicate the postnatal days (P) of injection (AM or PM). Pups were euthanized at P8 for analysis. (**F and G**) Representative flow cytometry plots of cell cycle speed analysis in iH2B-FT ECs (**F**) and quantification of fast-cycling iH2B-FT ECs (**G**) isolated from P8 livers of PBS (n=7), vehicle-treated BMP9/10ib (n=8), and palbociclib-treated BMP9/10ib (n=7) mice. Data in (**G**) represent mean ± SEM, one-way ANOVA with Tukey’s multiple comparisons test. ns, not significant; **P < 0.01, ***P < 0.001, ****P < 0.0001.

### Deregulations in restriction point activity occur in ECs of the HHT mice and patient skin telangiectasias

To gain insights into the mechanism of cell cycle deregulation in BMP9/10ib mice, liver ECs were isolated and assayed on a high-throughput ELISA for cell cycle proteins. Results showed that 21 proteins were upregulated by more than 10%, and among the top 10 upregulated proteins, six were mediators of the R point (E2F2, CDK4, Cyclin A1, E2F3, CDK2, and p27KIP1; **Fig. 3A**). qPCR analyses showed a significantly increased in *Cdk2* mRNA levels and a trend of increase in *Cdk4* expression in BMP9/10ib mouse liver ECs, compared to control ECs (**Fig. 3B**). Expression of mouse *Cdk6* (which is not part of the ELISA screen) was also significantly elevated in BMP9/10ib mouse liver ECs when measured by qPCR (**Fig. 3B**).

**Fig. 3.**
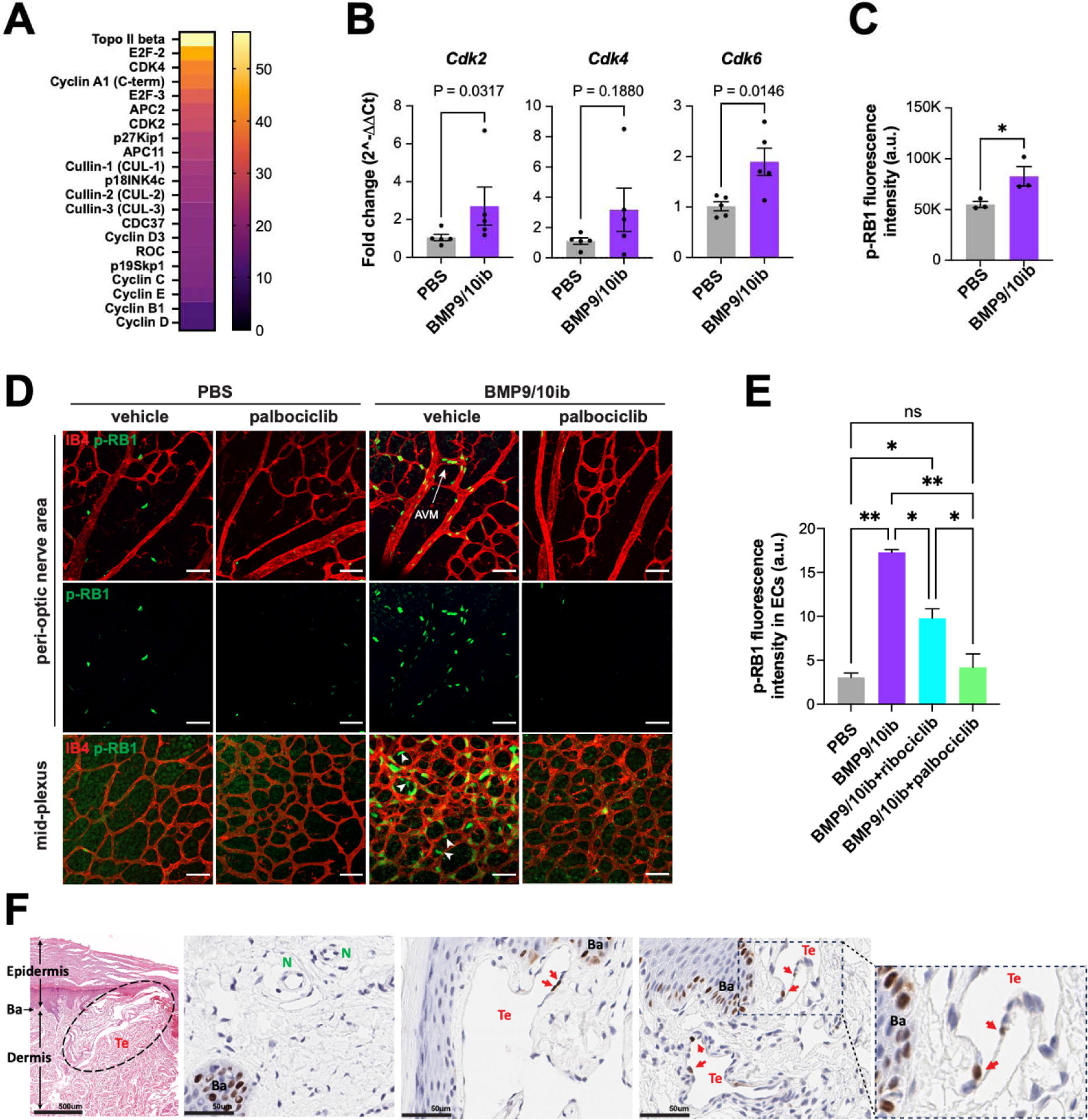
Restriction point activity in ECs of the HHT mice and patient skin telangiectasias. (**A**) Heatmap analysis of a high-throughput cell cycle protein ELISA of ECs isolated from livers of P6 BMP9/10ib mice vs. PBS control littermates. Data represent n=2 independent analyses (n represents one litter of pups combined for each condition; PBS, n=6 mice; BMP9/10ib, n=7 mice). (**B**) qPCR analysis of ECs isolated from livers of P6 PBS and BMP9/10ib mice (n=5/condition). Data represent mean ± SEM, Mann-Whitney test (*Cdk2*), and unpaired t-test (*Cdk4* and *Cdk6*). (**C**) Flow cytometry quantification of p-RB1 fluorescence intensity in ECs isolated from livers of P6 PBS and BMP9/10ib mice (n=3/condition). Data represent mean ± SEM, unpaired t-test. *P < 0.05. (**D and E**) Representative IF staining of p-RB1 (green) and IB4 (red) in the peri-optic nerve and mid-plexus regions (**D**) and corresponding quantification of the peri-optic nerve region (**E**) of retinas from vehicle-treated PBS (n=6), palbociclib-treated PBS (n=5), vehicle-treated BMP9/10ib (n=6), and palbociclib-treated BMP9/10ib (n=6) mice. Data in (**E**) represent individual retinas and mean ± SEM, one-way ANOVA with Tukey’s multiple comparisons test. ns, not significant; *P < 0.05, **P < 0.01. Scale bars in (**D**), 50 μm. (**F**) Representative H&E and IHC staining of p-RB1 in 4 μm skin sections of HHT2 patients. Te, telangiectasia; N, normal vessel; Ba, basilar cells.

Because RB1 is a master controller of the progression through the R point before S phase initiation, we next assessed its phosphorylation levels (phospho-Ser807/811-RB1, p-RB1) in BMP9/10ib mice by imaging flow cytometry (IFC) of isolated liver ECs and by immunofluorescence (IF) analysis of the retina. IFC showed an increase in the total median intensity of p-RB1 fluorescence in BMP9/10ib mouse liver ECs, compared to control ECs (**Fig. 3C**). In the retina, p-RB1 elevation was found in ECs of the AVMs and hypervascularized mid-plexus of the BMP9/10ib mice (**Figs. 3D, 3E**). Of note, increased p-RB1 immunoreactivity was also found in cells outside the endothelium in the hypervascularized mid-plexus of the BMP9/10ib retinas (white arrowheads), indicating that in addition to the ECs, other cell types proliferate at an increased rate in this retinal region. Thus, p-RB1 significantly increased in liver and retinal ECs of the BMP9/10ib mice.

We next assessed the p-RB1 level in skin telangiectasias of two HHT2 patients. Adult tissues express long-lived quiescent ECs that lack detectable proliferative activity. Single-EC transcriptomic analysis in mouse tissues has determined that only the liver and spleen contain measurable proliferation makers in about 1% of their ECs ^46^. We found that p-RB1 immunoreactivity was rare but consistently detected in ECs of the telangiectasias (4.0%, n = 742 ECs), while p-RB1^+^ ECs were found in significantly lower numbers in anatomically normal vessels in unaffected areas of the patient biopsies [0.7%, n = 453 ECs; P = 0.0004, Fisher’s exact test; OR (95% CI) = 5.510 (1.811-16.76), Woolf logit method] (**Fig. 3F**). Together, these results demonstrate that ALK1 signaling inhibition increased the expression of key R point mediators, including CDK4, CDK6, and CDK2 to promote p-RB1 elevation in vascular lesions of HHT mice and HHT patient skin biopsies.

### CDK4/6 kinase inhibitors, palbociclib and ribociclib, inhibit AVM pathology in HHT mice

Our data have shown that EC proliferation in the vascular defects of BMP9/10ib mice was accompanied by a G1/S checkpoint bypass, an acceleration of cell cycle speed, and an increase in R point activity. In this context, our observation that p-RB1 is potently elevated in ECs of the mouse AVMs and patient telangiectasias prompted us to determine whether pharmacological inhibition of CDK4/6 (the two main kinases targeting Ser807/811 on RB1) using the clinically approved drugs, palbociclib and ribociclib ^47^, could have beneficial effects in HHT mice. Palbociclib led to a nearly complete blockade of retinal AVM development (91% reduction on average, *P* < 0.0001, unpaired t-test), and ribociclib to a partial but significant reduction in AVM numbers (37%, *P* = 0.0207, unpaired t-test) in BMP9/10ib mice (**Figs. 4A-C**). Ribociclib also significantly reduced retinal AVM size (**Fig. 4D**), and palbociclib fully normalized both the increased vein diameter and hypervascularization caused by BMP9/10ib (**Figs. 4E, 4F**). Next, we assessed the anti-AVM effect of palbociclib in another HHT mouse model, the inducible conditional *Eng* knockout (*Eng*^iECKO^) mice ^39,48^. We found that palbociclib treatment significantly reduced AVM number and size (**Figs. 5A-C**), as well as hypervascularization of the plexus (**Figs. 5A, 5D**) in the retina of *Eng*^iECKO^ mice, compared to littermate controls treated with vehicle only.

**Fig. 4.**
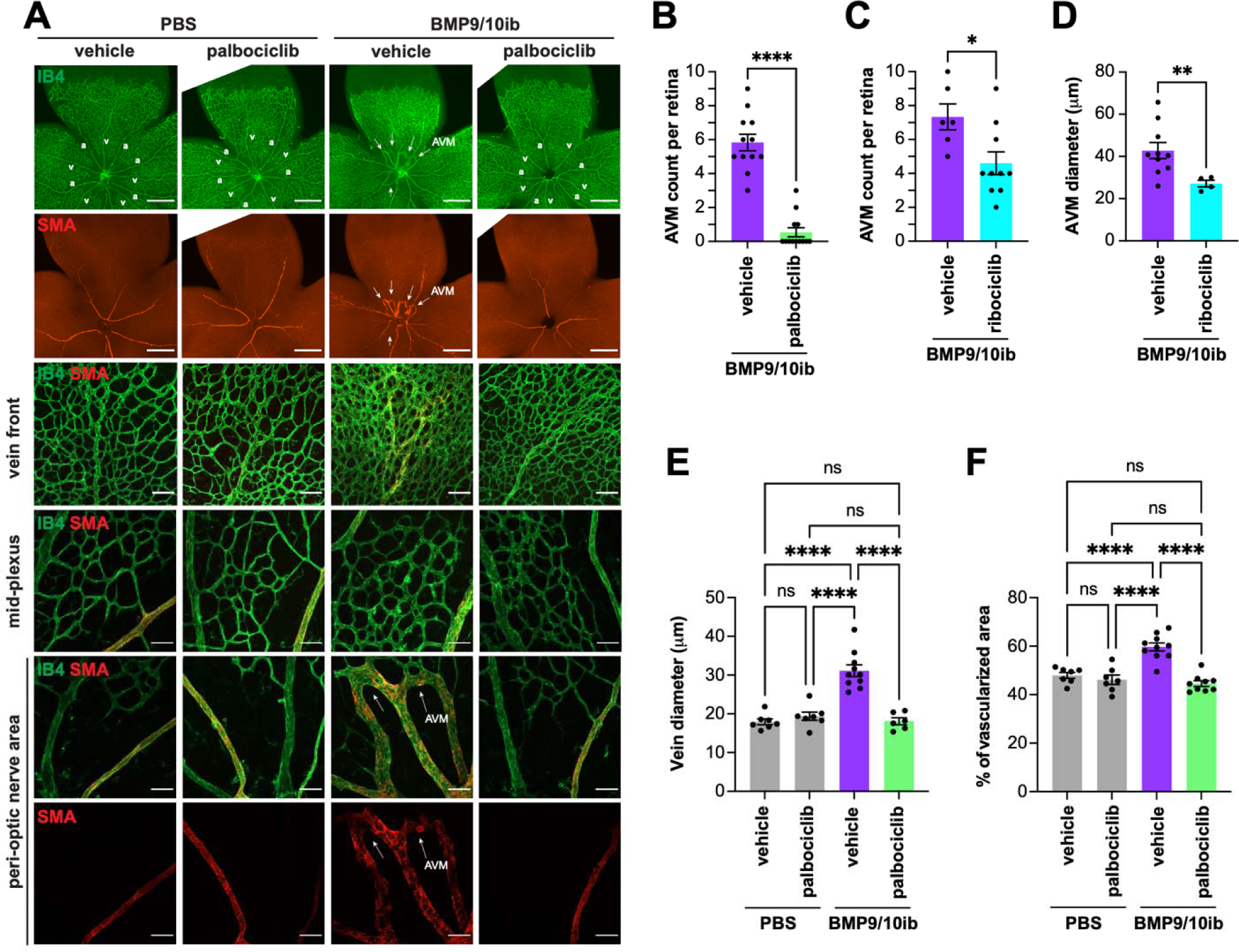
Effect of palbociclib and ribociclib on AVM pathology in BMP9/10ib mice. (**A**) Representative staining using IB4 (green) and of α-smooth muscle actin (SMA, red) in whole petals (first two rows), th vein front (third row), and peri-optic nerve (forth row) and mid-plexus (last two rows) regions of retinas from mice treated as indicated. Arrows denote AVMs; a, artery; v, vein. Scale bars, 1 mm (whole petal images), 100 μm (vein front), and 50 μm (mid-plexus and peri-optic nerve areas). (**B-F**) Scatter plots showing retinal AVM number (**B**), vein diameter (**E**), and mid-plexus vascular density (**F**) following palbociclib treatment, and retinal AVM number (**C**) and AVM diameter (**D**) following ribociclib treatment. Data represent individual retinas and mean ± SEM; vehicle-treated PBS (n=6 mice), palbociclib-treated PBS (n=7), vehicle-treated BMP9/10ib (n=10-12), and palbociclib-treated BMP9/10ib (n=9-13). Data represent individual retinas and mean ± SEM; unpaired t-test (**B-D**), one-way ANOVA with Tukey’s multiple comparisons test (**E and F**). *P 0.05, **P 0.01, ****P 0.0001.

**Fig. 5.**
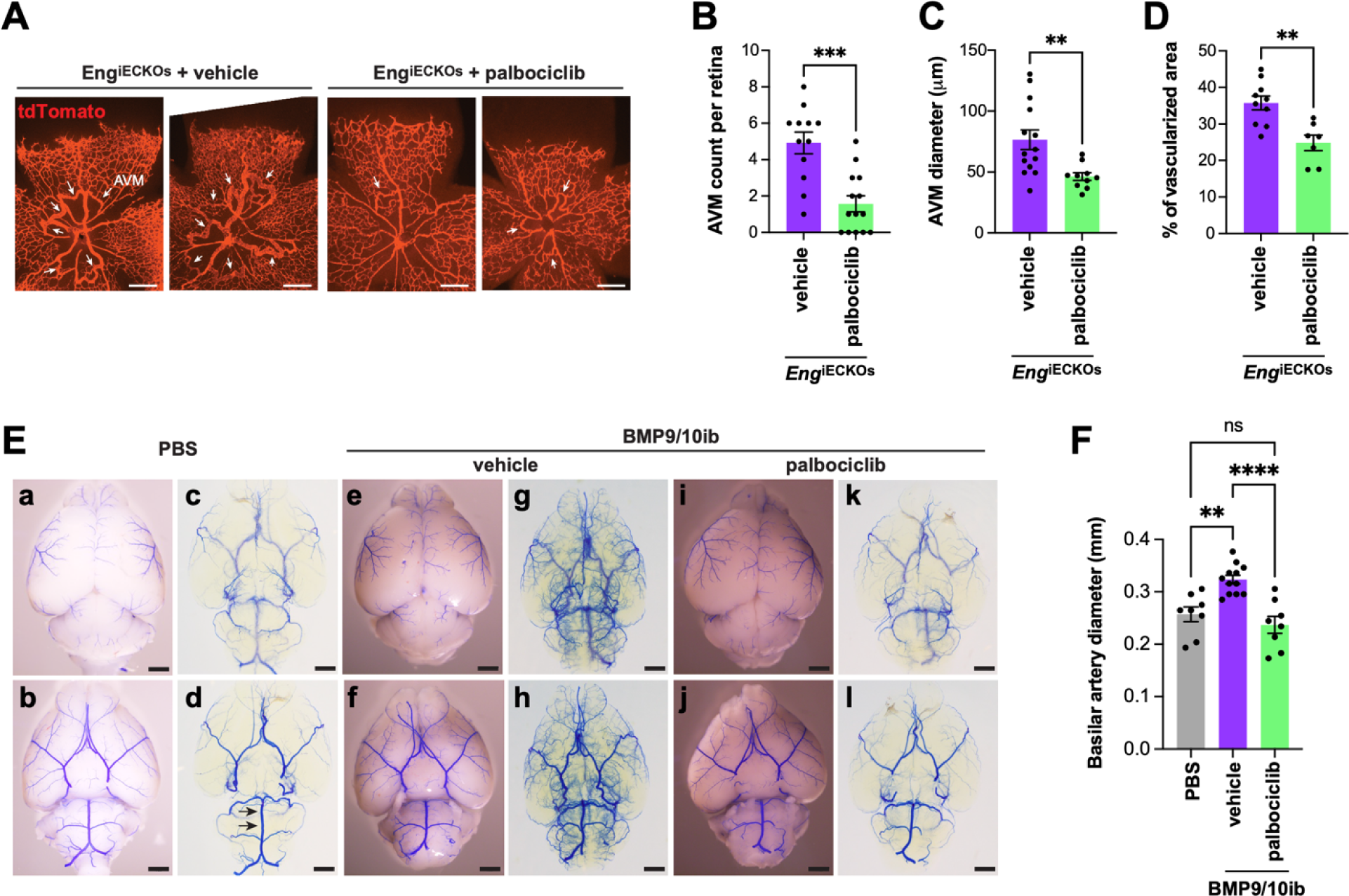
Effect of palbociclib on retinal and brain vascular pathologies in *Eng*^iECKO^ and BMP9/10ib mice. (**A**) Representative images of tdTomato-stained retinal vasculature from vehicle-treated and palbociclib-treated P6 *Eng*^iECKO^ mice. Arrows denote AVMs. Scale bars, 1 mm. (**B-D**) Scatter plot showing AVM number (**B**), AVM diameter (**C**), and mid-plexus vascular density (**D**) in retinas of mice treated as in (**A**). Data represent individual retinas and mean ± SEM; vehicle-treated *Eng*^iECKO^ (n=6 mice), palbociclib-treated *Eng*^iECKO^ (n=7); unpaired t-test. **P 0.01, ***P 0.001. (**E**) Representative bright field (**a, b, e, f, i, j**) and BABB-cleared (**c, d, g, h, k, l**) images of blue latex bead-perfused P8 PBS control and BMP9/10ib brains, treated or not (vehicle) with palbociclib. First row, dorsal views; second row, ventral views. Arrows in (**d**) denote the two positions where basilar artery (BA) diameter was measured. (**F**) Quantification of BA diameter across three groups: PBS (n=4), vehicle-treated BMP9/10ib (n=6), and palbociclib-treated BMP9/10ib (n=4) mice. Data represent two measurements per brain and mean ± SEM; one-way ANOVA with Tukey’s multiple-comparisons test. **P 0.01, ***P 0.001.

The effect of palbociclib on vascular pathology was investigated in the brain. To this end, blue latex beads were injected intracardially into the circulation to visualize arterial density and dilation. Due to their size, latex beads do not travel through the capillary system and remain in the arterial circulation. In the BMP9/10ib brains, compared to control brains, the dye revealed a hypervascularized and more tortuous deep arterial vasculature (visualized after BABB tissue clearing), a phenotype that could be prevented by palbociclib treatment (**Fig. 5E**). Diameter measurement of the basilar artery (BA) ^49^ revealed a significant vessel dilation in BMP9/10ib brains, compared to control brains; this effect was fully blocked by palbociclib (**Fig. 5F**). Thus, palbociclib efficiently blocked retinal AVMs and improved cerebral arterial hypervascularization and dilation in HHT mice.

### CDK4/6 inhibition normalizes endothelial cell cycle speed, proliferation, and cell number, and blocks endothelial p-RB1 elevation in HHT mice

Strikingly, treatment with palbociclib fully prevented the cell cycle acceleration observed in liver ECs of the iH2B-FT;BMP9/10ib mice (**Figs. 2F, 2G**). In addition, palbociclib significantly reduced EC proliferation (**Figs. 1B, 1E, 1F**) and the increase in EC numbers in the retina of BMP9/10ib mice (**Figs. 1B, 1C, 1D**). To confirm that the increase in p-RB1 observed in ECs of the BMP9/10ib retina was mechanistically linked to the anti-proliferative and anti-AVM effects of CDK4/6 inhibition, we asked whether palbociclib and ribociclib treatments affect p-RB1 level in the BMP9/10ib retina. In line with the effects of the two CDK4/6 inhibitors on AVM prevention (**Figs. 4B, 4C**), p-RB1 IF showed that treatments with palbociclib fully blocked the increase in p-RB1 detected in the AVMs and hyperproliferative mid-plexus, whereas ribociclib had a significant but only partial effect (**Figs. 3D, 3E**). Thus, the reduction in AVM pathology by CDK4/6 inhibitors is accompanied by a normalization of endothelial cell cycle speed, proliferation, and cell number, and a reduction of endothelial p-RB1 elevation in HHT mice.

### Endothelial *Cdk6* deletion blocks AVM development in HHT mice

Because p-RB1 elevation was also found in cells outside the hyperproliferating vessels of the BMP9/10ib retina (**Fig. 3D**) and because CDK4/6 inhibitors can also block the proliferation of non-endothelial cells that could contribute to the studied vascular defects, we sought to determine whether EC-specific *Cdk6* deficiency is sufficient to alter vascular pathology development in HHT mice. We focus on CDK6 because previous work in cancer has highlighted its specific contribution to tumor angiogenesis ^50^. BMP9/10ib mice deficient for endothelial *Cdk6* were generated, and retinas were analyzed for the presence of AVMs and hypervascularization. Inducible EC-specific *Cdk6* KO (*Cdk6*^iECKO^) mice were obtained by crossing *Cdk6*^f/f^ mice with *Cdh5*-*Cre*^ERT2^ mice. Gene deletion was induced by administering tamoxifen at P1 and P2, while BMP9/10ib was induced as before at P3. *Cdk6*^iECKO^;BMP9/10ib mice were significantly protected from retinal AVM, vein dilation, and hypervascularization pathologies compared to their littermate controls (*Cre*-negative Cdk6^f/f^;BMP9/10ib mice; **Figs. 6A-D**). Thus, *Cdk6* deletion is sufficient to block vascular pathology in HHT mice.

**Fig. 6.**
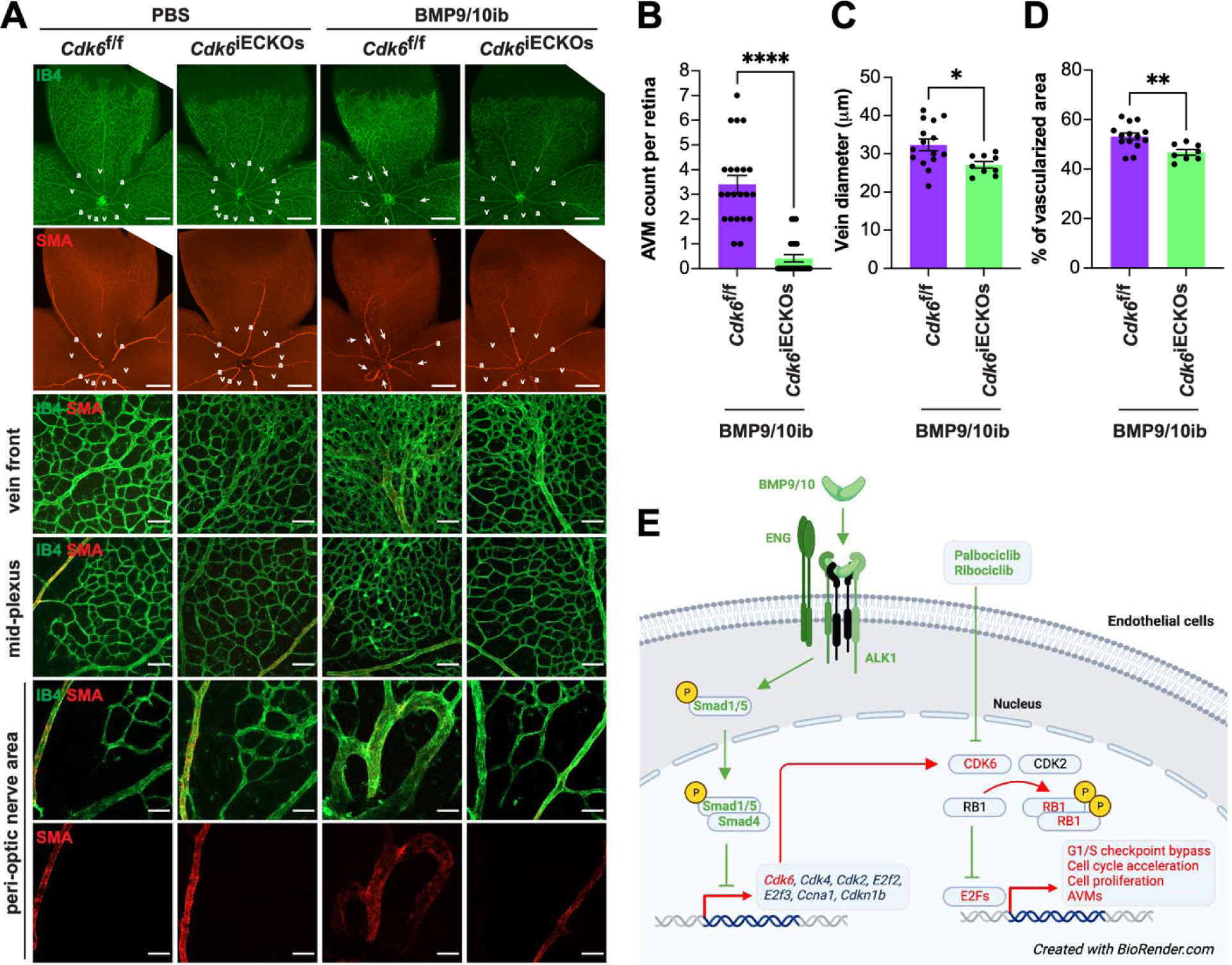
Effect of *Cdk6* deletion on AVM pathology in BMP9/10ib mice. (**A**) Representative staining using IB4 (green) and of SMA (red) in whole petals (first two rows), the vein front (third row), and peri-optic nerve (forth row) and mid-plexus (last two rows) regions of retinas from *Cdk6*^f/f^ controls and *Cdk6*^iECKO^ mice challenged or not (PBS) with BMP9/10ib. Arrows denote AVMs; a, artery; v, vein. Scale bars, 1 mm (whole petal images), 100 μm (vein front), and 50 μm (mid-plexus and peri-optic nerve areas). (**B-D**) Scatter plots showing retinal AVM number (**B**), vein diameter (**C**), and mid-plexus vascular density (**D**) in *Cdk6*^f/f^;BMP9/10ib controls (n=11-14) and *Cdk6*^iECKO^;BMP9/10ib (n=8-12) mice. Data represent individual retinas and mean ± SEM, unpaired t-test. **P 0.01, ***P 0.001. (**D**) Schematic illustration of the proposed mechanism of control of the cell cycle in ECs by ALK1 signaling and its relevance for HHT pathogenesis.

## DISCUSSION

EC accumulation in AVMs is a central hallmark of HHT. Previous data have provided compelling evidence that ALK1 signaling controls the cell cycle by suppressing G1 to S/G2/M transition ^28,29^. Using a unique reporter mouse model of cell cycle speed assessment, we report here for the first time that, in addition to undergoing a G1/S checkpoint bypass (**Fig. 2**), live ECs of BMP9/10ib mice exhibited a significant acceleration of their cell cycle (**Fig. 2**), demonstrating the presence of a complete change of cycling behavior in these cells. We further found that p-RB1 increased in ECs of BMP9/10ib mouse AVMs and HHT patient skin telangiectasias. Our observation that p-RB1 and, more generally, R point activity (*via* the increased levels of key R point mediators) were elevated (**Fig. 3**), offered the opportunity to intervene pharmacologically with clinically approved CDK4/6 inhibitors. Indeed, a central function of cyclin D-CDK4/6 in driving cell cycle progression is to phosphorylate and inactivate RB1 ^26^. Our data revealed that palbociclib demonstrated efficacy at slowing down endothelial cell cycle speed and proliferation (**Figs. 1 and 2**), as well as blocked AVM development in BMP9/10ib and *Eng*^iECKO^ mice (**Figs. 4 and 5**). Together these results reveal a mechanism of control of EC proliferation required for HHT pathogenesis, in which ALK1 signaling LOF increased endothelial cell cycle speed and progression *via* the overexpression of key mediators of the R point (including CDK6 and CDK2) to promote RB1 inactivation and S phase initiation (**Fig. 2**). Consequently, CDK4/6 inhibition using palbociclib, which precisely acts at the level of RB1, demonstrated robust anti-AVM properties.

The aberrant cell cycling and proliferation phenotypes of the AVM ECs and the anti-AVM activity associated with palbociclib (and, to a certain extent, with ribociclib) draw compelling parallels between AVM and tumor development. Blocking tumoral cell division is a common approach to cancer treatment, and deregulations of the cyclin D-CDK4/6-RB1-E2F pathway are found in about 40% of all human tumors ^51,52^. In addition, cyclin D-CDK4/6 activity is controlled by mitogenic pathways, such as PI3K and mTOR, which are also frequently deregulated in cancer and are abnormally elevated in AVM ECs ^20,29,35^. In cerebral cavernous malformations (CCMs), a vascular malformation syndrome of the central nervous system, the comparison between excessive vessel growth and cancer has already been made ^53^. The authors suggested that the LOF of constitutive vascular suppressor genes works with a vascular oncogene (identified in the mTOR signaling pathway) to drive CCMs. We propose that a similar cancer-like mechanism is at play during HHT AVM development, where LOF of HHT genes lowers the response threshold of R point activity to mitogenic signals during pro-angiogenic stimulation of the ECs. Knowing that concordant data have shown that one of these signals is PI3K-mTOR, combination therapies targeting both CDK4/6 and PI3K-mTOR pathways could be explored. In this context, we and others have identified clinically approved mTOR inhibitor sirolimus as a potential therapy for HHT ^20^, CCMs ^53^, and vascular anomalies ^54^. Furthermore, it was reported that non-resident/circulating ECs carrying HHT mutations could contribute to lesion development, suggesting the presence of a mechanism of disease spreading that could also be compared to cancer pathogenesis. Indeed, transplantation of *Alk1*-deficient bone marrow cells in wild mice led to brain AVMs upon VEGFA stimulation ^55^, and liver transplantation of HHT patients from healthy donors resulted in the recurrence of hepatic AVM disease in the recipient patients ^56^. In sum, we propose that the current work describes a defective cell cycle pathway with cancer-like properties involved in AVM development and identifies anti-cancer CDK4/6 inhibitors as potential therapeutics for HHT.

We found that EC-specific *Cdk6* KO was sufficient to protect HHT mice from AVM pathology (**Fig. 6**), suggesting that CDK6 is uniquely involved in the cell cycle defects leading to EC proliferation and AVM development. Palbociclib, ribociclib, and abemaciclib are the currently clinically approved CDK4/6 kinase inhibitors prescribed for advanced or metastatic hormone receptor-positive breast cancer ^57^. Although these drugs are usually well tolerated and thus have promising translational potential in HHT, they are associated with some toxicities, such as anemia consecutive to hematopoietic defects ^25^. CDK4 and CDK6 are paralogues with overlapping and independent functions. Constitutive CDK4/6 double KO is embryonic lethal, but CDK4 or CDK6 single KO mice are viable and have distinct phenotypes. The CDK6 KOs are generally healthier despite showing alteration of the hematopoietic system. This study showed no apparent toxicities in mouse neonates associated with CDK4/6 inhibitor treatment or CDK6 KO. Nevertheless, it will be interesting in future studies to investigate and compare the potential effects in the long term of these interventions on the hematopoietic system, knowing that some HHT models are predisposed to hemorrhage and develop anemia. In addition, it was reported that CDK6 has kinase-independent pro-angiogenic functions *via* the direct transcriptional control of VEGFA ^50^. In this context, it will also be interesting to assess and compare the effects of the CDK4/6 inhibitors and CDK6 KO on VEGF signaling and determine whether inhibiting this pathway contributes to the anti-AVM properties of CDK6 deletion.

Our results align with studies showing that G1 arrest is required for arterial specification ^58^ and that cell cycle arrest using palbociclib could prevent arteriovenous development defects in *Cx37*-null mice ^59^. In BMP9/10ib mice, AVM pathology is accompanied by arteriovenous specification defects and muscularization of the veins and AVMs by SMCs (SMA staining, **Figs. 4A and 6A**). We found that both CDK4/6 inhibitor treatment and CDK6 KO efficiently rescued arteriovenous specification in BMP9/10ib mice (**Figs. 4A and 6A**). How arteriovenous specification defects and AVMs crosstalk during HHT pathogenesis remains a knowledge gap; because this work demonstrates that cell cycle acceleration and CDK6 activity are involved in both processes, it paves the way to future investigations aimed at delineating the importance of this crosstalk in AVM pathogenesis.

In conclusion, our work reveals a mechanism of endothelial cell cycle deregulation involved in excessive EC proliferation and AVM development, in which cell cycle acceleration and CDK6 activity play pivotal roles. We further identify clinically approved CDK4/6 inhibitors as potential disease-modifying therapies for HHT.

## MATERIALS AND METHODS

### Mice and treatments

All animal procedures were performed in accordance with protocols approved by The Feinstein Institutes for Medical Research Institutional Animal Care and Use Committees (IACUC) and conformed to the NIH Guide for the Care and Use of Laboratory Animals and ARRIVE guidelines. C57BL/6J mice from The Jackson Laboratory were used for the BMP9/10ib studies. BMP9 (25 mg/kg; MAB3209, R&D System) and BMP10 (50 mg/kg; MAB2926, R&D System) were injected intraperitoneally (i.p.) at P3 and P4. Mice were then sacrificed at P6 or P8, depending on the analysis. Retinas, livers, and brains were collected unless mice were used for latex bead injection. iH2B-FT knock-in mice, which were generated by targeting the H2B-FT coding sequence into the *HPRT* locus under a doxycycline-inducible promoter ^45^, were obtained from the MMRRC via The Jackson Laboratory (#066960) and crossed with R26-M2rtTA mice (The Jackson Laboratory, #006965). To induce H2B-FT expression, iH2B-FT mice were injected i.p. daily for three consecutive days at P5-P7 with 200 mg/kg doxycycline hyclate (Sigma Aldrich, D9891). iH2B-FT mice were then analyzed at P8. To determine the baseline fluorescence for H2B-FT expression analysis, colorless R26-M2rtTA mice were used as negative controls. *Cdk6*^f/f^ mice (C57BL/6N-Cdk6tm1c(EUCOMM)Wtsi/Tcp mouse line, Canadian Mouse Mutant Repository) were generated with C57BL/6N-Cdk6tm1a(EUCOMM)Wtsi/Tcp line made from EUCOMM ES cells ^60^ at the Toronto Center for Phenogenomics, Canada. *Eng*^f/f^ ^39,48^ and *Cdk6*^f/f^ mice were crossed with *Cdh5*-Cre^ERT2^ mice [Taconic, #13073; ^61^] to generate EC-specific, tamoxifen-inducible KO mice (*Eng*^iECKO^ and *Cdk6*^iECKO^). *Eng*^iECKO^ mice were also crossed with *Rosa26*-tdTomato reporter mice (Ai14, #007914, The Jackson Laboratory) to assess recombination efficiency and visualize retinal vascular pathology. Gene deletion was induced in *Eng*^iECKO^ mice by i.p. injection of 25 μg Z-(4)-hydroxytamoxifen (Sigma, H7904) at P1 and P2, and in *Cdk6*^iECKO^ mice by intragastric administration of 300 μg tamoxifen (Sigma, T5648) at P1 and P2. *Cdk6*^iECKO^ mice were injected with BMP9/10 blocking antibodies (as described above) to generate *Cdk6*^iECKO^;BMP9/10ib mice. *Cdk6* deletion was verified in isolated liver ECs (see below for Methods) from one litter by TaqMan qPCR (*Cdk6* probe assay ID# Mm01311342_m1 and housekeeping *Gapdh* probe assay ID# Mm99999915_g1, ThermoFisher). 66.9% reduction in *Cdk6* expression was measured in *Cdk6*^iECKO^ (n=4) compared to *Cdk6*^f/f^ (n=2) mice. The sex of the pups was not determined, as HHT affects males and females equally. Mice were injected i.p. at P3, P4, and P5 with palbociclib isethionate (MedChemExpress, HY-A0065) in saline (100 mg/kg) or with ribociclib hydrochloride (MedChemExpress, HY-15777A) in 10% DMSO, 20% SBE-β-CD (MedChemExpress, HY-17031) in saline (100 mg/kg). Mice were maintained in regular housing conditions and were allowed free access to water and a maintenance diet.

### Human skin sample analyses

Skin telangiectasias were resected using a 3 mm punch biopsy after local anesthesia (1% xylocaine with epinephrine) with a standard aseptic technique. Samples were immediately formalin-fixed (10% formalin) and then shipped at room temperature. The telangiectasias were resected as part of an exploratory aim of the clinical trial “Doxycycline for Hereditary Hemorrhagic Telangiectasia” (NCT03397004), which was a negative clinic trial ^62^. The clinical trial protocol was approved by the Research Ethics Board at St. Michael’s Hospital (REB) and was funded by Department of Defense grant W81XWH-17-1-0429. Biopsies were processed, sectioned (4 μm), and stained with hematoxylin/eosin (H&E) and anti-p-RB1 (Ser807/811) antibody (Cell Signaling Technology, # 4277) at HistoWiz.

### Retinal immunofluorescence (IF)

Retinas were processed as described before ^18–20^. Briefly, after sacrifice, mouse eyes were dissected and immersed completely in ice-cold 4% paraformaldehyde (PFA; Sigma Aldrich, 158127) in phosphate-buffered saline (PBS) for 30 min. Eyes were then transferred to ice-cold PBS for dissection. The cornea, lens, and pigmented layers were removed from the retinas and four incisions were made with Vannas spring scissors (Fine Science Tools, 91500-09) for flat mounting. PBS was removed, and retinas were immersed in 100% ice-cold methanol for storage or used immediately for IF. Retinas were rinsed with PBS and immersed in blocking solution [10% heat-inactivated serum (HIS)/0.2% tween PBS (PBST)] for 1h at room temperature. Retinas were then stained at 4°C overnight with isolectin GS1-IB4 conjugated to Alexa Fluor 488 or Alexa Fluor 568 (ThermoFisher Scientific, I21411 and I21412; 1:300 in blocking solution) and with anti-alpha smooth muscle actin-Cy3 antibody (Sigma Aldrich, C6198; 1:100 dilution in blocking solution) and anti-ERG (Abcam, ab92513; 1:250 in blocking solution). Retinas were then washed three times in 2% HIS/PBST. If only pre-conjugated probes were used, retinas were washed once more in PBS and then mounted on slides using anti-fade mountant (ThermoFisher, P36965) and sealed with nail polish. For ERG IF, goat-anti-Rabbit IgG (H+L) Pacific Blue secondary antibody was used at a dilution of 1:750 in 2% HIS/PBST by immersion at room temperature for 1 h. Retinas were then washed three times in 2% HIS/PBST and once in PBS before mounting. For p-RB1 IF, anti-phospho-RB1 (Ser807/811) antibody (Cell Signaling Technology, #8516; 1:200 in blocking solution) and goat anti-rabbit IgG, Alexa Fluor 488 tyramide SuperBoost kit (ThermoFisher, B40922) were used, as per manufacturer recommendation. For EdU detection, we used the Click-iT EdU Cell Proliferation kit, conjugated to Alexa Fluor 647 (ThermoFisher, C10340), as per manufacturer recommendation. Retinas were imaged on a Zeiss LSM900 confocal microscope for 10X and 20X magnifications and on a ThermoFisher EVOS M7000 microscope for 2X magnifications.

### EC isolation

Liver ECs were collected from fresh liver tissue for flow cytometry. Livers were collected and placed in ice-cold DMEM. DMEM was removed and replaced with a warm collagenase/dipase solution, and the tissues were homogenized by mechanical separation with a sterile blade. The homogenates were then incubated for 15 min at 37°C and triturated with a cannula 12-15 times. Homogenates were incubated for another 15 min and then triturated again with a cannula 12-15 times before filtering with a 70 μm strainer (ThermoFisher, 437150). An equal volume of 5% FBS in DMEM was then added to the homogenates to neutralize digestion. Samples were centrifuged at 400xg for 15 min and resuspended in 5% BSA, 2 mM EDTA in PBS. To isolate the ECs, CD31 microbeads were added to the cell suspensions, per the manufacturer’s recommendation (Milltenyi Biotec, 130-097-418). Suspensions were incubated at 4°C for 15 min. MS columns and the octoMACS separator were then used to isolate CD31^+^ liver cells, per manufacturer recommendation (Milltenyi Biotec, 130-042-201 and 130-042-109).

### Flow cytometry

Cells were labeled with CD31-FITC antibody (0.5 μg/sample) to validate EC isolation (BD-Pharmigen, 558738) and incubated on ice for 30 min. Cells were fixed in ice-cold 1% PFA for 10 min. For EdU cell cycle phase determination, we used the Click-iT EdU Pacific Blue flow cytometry assay kit, per manufacturer recommendation (ThermoFisher, C10418). Cells were analyzed on a BD-FACSymphony Flow Cytometer. For iH2B-FT cell flow analysis, only live cells were processed for cell cycle speed determination to avoid photoconversion ^45^. For imaging flow cytometry (IFC), cells were labeled with CD31-FITC and fixed, as described above. Cells were then permeabilized with saponin buffer (0.1% saponin, 0.5% BSA in PBS) and stained for intracellular p-RB1 [Cell Signaling Technology, # 8156; 1:500 in flow buffer (0.5% BSA in PBS)] for 1h. Cells were centrifuged at 1200xg at 4°C and washed twice in ice-cold PBS. Cells were stained with Alexa Fluor 532 secondary antibody (ThermoFisher, A20182; 1:1000 in flow buffer) and centrifuged at 1200xg at 4°C and washed three times in ice-cold PBS before a final resuspension of approximately 100,000 cells/tube in DAPI-supplemented PBS for final analysis. Cells were analyzed on an Amnis ImageStream cytometer. Analyses were performed using IDEAS software and the bright detail intensity feature was used.

### Proteome array and qPCR

Liver ECs were isolated as described above from BMP9/10ib mice and their littermate controls. A high-throughput ELISA was performed per manufacturer’s recommendation (Full Moon Biosystems, ACC058). Array scans were analyzed using ImageJ *via* the MicroArray Profiler Plugin (Optinav). TaqMan qPCR was performed on liver ECs using *Cdk2* probe assay ID# Mm00443947_m1, *Cdk4* probe assay ID# Mm00726334_s1, *Cdk6* probe assay ID# Mm01311342_m1, and housekeeping *Gapdh* probe assay ID# Mm99999915_g1 (ThermoFisher).

### Brain latex bead injections and BABB clearing

P8 pups were processed for cardiac perfusion after euthanasia, following procedures described before ^18,20^. Briefly, the left ventricles were injected manually with 600 μL of blue latex beads (470024-612, Ward’s Science) using an insulin syringe, and the right atrium was opened to drain the blood. Brains were dissected, fixed in 4% PFA, and washed in PBS. Blocks were dehydrated in methanol series and cleared with organic solvent [benzyl alcohol/benzyl benzoate (BABB), 1:2; Sigma]. Images of the whole brain were acquired using an Olympus SZX7 stereomicroscope attached to an Olympus DP27 camera.

### ImageJ Analyses

For vascular occupancy measurements, we used the plugin on ImageJ to analyze skeletonized images of IB4-stained retinas. An adjusted model based on the vessel density plugin determined the percentage of area occupied by the IB4^+^ signal ^63^. For vein and AVM diameter measurements, vessels within 500 μm of the optic nerve were used. All vessels were measured by their largest diameter within the 500 μm range. Measurements were calibrated and taken on ImageJ. IF intensity measurements were conducted on ImageJ using the channel intensity tool specific to the channel of interest, depending on the fluorophore.

### Statistical analysis

Data were analyzed with GraphPad Prism 10 (GraphPad Software, San Diego, CA, USA; www.graphpad.com). All statistical details can be found in the figures and figure legends. P <

0.05 was considered statistically significant. All values are expressed as means ± SEM.

## Acknowledgments

*Eng*^iECKO^ mice were kindly provided by Dr. Helen M. Arthur (University of Newcastle, Newcastle, UK) through the assistance of Dr. Stryder M. Meadow (Tulane University, New Orleans, LA, USA).

## Author contributions

Conceptualization: SD, PM. Methodology: SD, HZ, YT, ZW, SR, ANK, LB, MEF, PM. Writing of original draft: SD, PM; review & editing of the manuscript: LB, MEF.

## Data and materials availability

All data are available in the main text.

## REFERENCES

1. Guttmacher, A. E., Marchuk, D. A. & White, R. I. Hereditary hemorrhagic telangiectasia. N. Engl. J. Med. 333, 918–924 (1995).

2. Brinjikji, W., Iyer, V. N., Wood, C. P. & Lanzino, G. Prevalence and characteristics of brain arteriovenous malformations in hereditary hemorrhagic telangiectasia: a systematic review and meta-analysis. J. Neurosurg. 127, 302–310 (2017).

3. Faughnan, M. E. et al. Second International Guidelines for the Diagnosis and Management of Hereditary Hemorrhagic Telangiectasia. Ann. Intern. Med. (2020) doi:10.7326/M20-1443.

4. Shovlin, C. L. Hereditary haemorrhagic telangiectasia: pathophysiology, diagnosis and treatment. Blood Rev. 24, 203–219 (2010).

5. Dupuis-Girod, S., Bailly, S. & Plauchu, H. Hereditary hemorrhagic telangiectasia: from molecular biology to patient care. J. Thromb. Haemost. 8, 1447–1456 (2010).

6. Snodgrass, R. O., Chico, T. J. A. & Arthur, H. M. Hereditary Haemorrhagic Telangiectasia, an Inherited Vascular Disorder in Need of Improved Evidence-Based Pharmaceutical Interventions. Genes 12, (2021).

7. Robert, F., Desroches-Castan, A., Bailly, S., Dupuis-Girod, S. & Feige, J.-J. Future treatments for hereditary hemorrhagic telangiectasia. Orphanet J. Rare Dis. 15, 4 (2020).

8. McAllister, K. A. et al. Endoglin, a TGF-beta binding protein of endothelial cells, is the gene for hereditary haemorrhagic telangiectasia type 1. Nat. Genet. 8, 345–351 (1994).

9. Johnson, D. W. et al. Mutations in the activin receptor-like kinase 1 gene in hereditary haemorrhagic telangiectasia type 2. Nat. Genet. 13, 189–195 (1996).

10. Gallione, C. J. et al. A combined syndrome of juvenile polyposis and hereditary haemorrhagic telangiectasia associated with mutations in MADH4 (SMAD4). Lancet 363, 852–859 (2004).

11. Kim, S. K., Henen, M. A. & Hinck, A. P. Structural biology of betaglycan and endoglin, membrane-bound co-receptors of the TGF-beta family. Exp. Biol. Med. 244, 1547–1558 (2019).

12. Desroches-Castan, A., Tillet, E., Bouvard, C. & Bailly, S. BMP9 and BMP10: Two close vascular quiescence partners that stand out. Dev. Dyn. 251, 178–197 (2022).

13. Arthur, H. M. & Roman, B. L. An update on preclinical models of hereditary haemorrhagic telangiectasia: Insights into disease mechanisms. Front. Med. 9, 973964 (2022).

14. Ricard, N., Bailly, S., Guignabert, C. & Simons, M. The quiescent endothelium: signalling pathways regulating organ-specific endothelial normalcy. Nat. Rev. Cardiol. 18, 565–580 (2021).

15. Snellings, D. A. et al. Somatic Mutations in Vascular Malformations of Hereditary Hemorrhagic Telangiectasia Result in Bi-allelic Loss of ENG or ACVRL1. Am. J. Hum. Genet. (2019) doi:10.1016/j.ajhg.2019.09.010.

16. Al-Samkari, H. et al. An international, multicenter study of intravenous bevacizumab for bleeding in hereditary hemorrhagic telangiectasia: the InHIBIT-Bleed study. Haematologica 106, 2161–2169 (2021).

17. Dupuis-Girod, S. et al. European Reference Network for Rare Vascular Diseases (VASCERN): When and how to use intravenous bevacizumab in Hereditary Haemorrhagic Telangiectasia (HHT)? Eur. J. Med. Genet. 65, 104575 (2022).

18. Ruiz, S. et al. A mouse model of hereditary hemorrhagic telangiectasia generated by transmammary-delivered immunoblocking of BMP9 and BMP10. Sci. Rep. 5, 37366 (2016).

19. Ruiz, S. et al. Tacrolimus rescues the signaling and gene expression signature of endothelial ALK1 loss-of-function and improves HHT vascular pathology. Hum. Mol. Genet. 26, 4786– 4798 (2017).

20. Ruiz, S. et al. Correcting Smad1/5/8, mTOR, and VEGFR2 treats pathology in hereditary hemorrhagic telangiectasia models. J. Clin. Invest. 130, 942–957 (2020).

21. Roman, B. L. & Hinck, A. P. ALK1 signaling in development and disease: new paradigms. Cell. Mol. Life Sci. 74, 4539–4560 (2017).

22. Baeyens, N., Bandyopadhyay, C., Coon, B. G., Yun, S. & Schwartz, M. A. Endothelial fluid shear stress sensing in vascular health and disease. J. Clin. Invest. 126, 821–828 (2016).

23. Augustin, H. G. & Koh, G. Y. Organotypic vasculature: From descriptive heterogeneity to functional pathophysiology. Science 357, (2017).

24. Potente, M. & Mäkinen, T. Vascular heterogeneity and specialization in development and disease. Nat. Rev. Mol. Cell Biol. 18, 477–494 (2017).

25. Fassl, A., Geng, Y. & Sicinski, P. CDK4 and CDK6 kinases: From basic science to cancer therapy. Science 375, eabc1495 (2022).

26. Goel, S., Bergholz, J. S. & Zhao, J. J. Targeting CDK4 and CDK6 in cancer. Nat. Rev. Cancer 22, 356–372 (2022).

27. Fernandez-L, A. et al. Gene expression fingerprinting for human hereditary hemorrhagic telangiectasia. Hum. Mol. Genet. 16, 1515–1533 (2007).

28. Lamouille, S., Mallet, C., Feige, J.-J. & Bailly, S. Activin receptor-like kinase 1 is implicated in the maturation phase of angiogenesis. Blood 100, 4495–4501 (2002).

29. Ola, R. et al. PI3 kinase inhibition improves vascular malformations in mouse models of hereditary haemorrhagic telangiectasia. Nat. Commun. 7, 13650 (2016).

30. Lin, K. et al. Molecular mechanism of endothelial growth arrest by laminar shear stress. Proc. Natl. Acad. Sci. U. S. A. 97, 9385–9389 (2000).

31. Han, Z. et al. Aryl hydrocarbon receptor mediates laminar fluid shear stress-induced CYP1A1 activation and cell cycle arrest in vascular endothelial cells. Cardiovasc. Res. 77, 809–818 (2008).

32. Fang, J. S. et al. Shear-induced Notch-Cx37-p27 axis arrests endothelial cell cycle to enable arterial specification. Nat. Commun. 8, 2149 (2017).

33. Baeyens, N. et al. Defective fluid shear stress mechanotransduction mediates hereditary hemorrhagic telangiectasia. J. Cell Biol. 214, 807–816 (2016).

34. Cuyàs, E., Corominas-Faja, B., Joven, J. & Menendez, J. A. Cell Cycle Regulation by the Nutrient-Sensing Mammalian Target of Rapamycin (mTOR) Pathway. in Cell Cycle Control: Mechanisms and Protocols (eds. Noguchi, E. & Gadaleta, M. C.) 113–144 (Springer New York, 2014).

35. Alsina-Sanchís, E. et al. ALK1 Loss Results in Vascular Hyperplasia in Mice and Humans Through PI3K Activation. Arterioscler. Thromb. Vasc. Biol. 38, 1216–1229 (2018).

36. David, L. et al. Bone morphogenetic protein-9 is a circulating vascular quiescence factor. Circ. Res. 102, 914–922 (2008).

37. Scharpfenecker, M. et al. BMP-9 signals via ALK1 and inhibits bFGF-induced endothelial cell proliferation and VEGF-stimulated angiogenesis. J. Cell Sci. 120, 964–972 (2007).

38. Tual-Chalot, S. et al. Endothelial depletion of Acvrl1 in mice leads to arteriovenous malformations associated with reduced endoglin expression. PLoS One 9, e98646 (2014).

39. Mahmoud, M. et al. Pathogenesis of arteriovenous malformations in the absence of endoglin. Circ. Res. 106, 1425–1433 (2010).

40. Crist, A. M., Lee, A. R., Patel, N. R., Westhoff, D. E. & Meadows, S. M. Vascular deficiency of Smad4 causes arteriovenous malformations: a mouse model of Hereditary Hemorrhagic Telangiectasia. Angiogenesis 21, 363–380 (2018).

41. Jin, Y. et al. Endoglin prevents vascular malformation by regulating flow-induced cell migration and specification through VEGFR2 signalling. Nat. Cell Biol. 19, 639–652 (2017).

42. Ola, R. et al. SMAD4 Prevents Flow Induced Arterial-Venous Malformations by Inhibiting Casein Kinase 2. Circulation 138, 2379–2394 (2018).

43. Choi, H. et al. BMP10 functions independently from BMP9 for the development of a proper arteriovenous network. Angiogenesis (2022) doi:10.1007/s10456-022-09859-0.

44. Hwang, Y. et al. Global increase in replication fork speed during a p57KIP2-regulated erythroid cell fate switch. Sci Adv 3, e1700298 (2017).

45. Eastman, A. E. et al. Resolving Cell Cycle Speed in One Snapshot with a Live-Cell Fluorescent Reporter. Cell Rep. 31, 107804 (2020).

46. Kalucka, J. et al. Single-Cell Transcriptome Atlas of Murine Endothelial Cells. Cell 180, 764–779.e20 (2020).

47. O’Leary, B., Finn, R. S. & Turner, N. C. Treating cancer with selective CDK4/6 inhibitors. Nat. Rev. Clin. Oncol. 13, 417–430 (2016).

48. Allinson, K. R., Carvalho, R. L. C., van den Brink, S., Mummery, C. L. & Arthur, H. M. Generation of a floxed allele of the mouse Endoglin gene. Genesis 45, 391–395 (2007).

49. Zhou, X. et al. ANG2 Blockade Diminishes Proangiogenic Cerebrovascular Defects Associated With Models of Hereditary Hemorrhagic Telangiectasia. Arterioscler. Thromb. Vasc. Biol. (2023) doi:10.1161/ATVBAHA.123.319385.

50. Kollmann, K. et al. A kinase-independent function of CDK6 links the cell cycle to tumor angiogenesis. Cancer Cell 24, 167–181 (2013).

51. Sherr, C. J., Beach, D. & Shapiro, G. I. Targeting CDK4 and CDK6: From Discovery to Therapy. Cancer Discov. 6, 353–367 (2016).

52. Bonelli, M., La Monica, S., Fumarola, C. & Alfieri, R. Multiple effects of CDK4/6 inhibition in cancer: From cell cycle arrest to immunomodulation. Biochem. Pharmacol. 170, 113676 (2019).

53. Ren, A. A. et al. PIK3CA and CCM mutations fuel cavernomas through a cancer-like mechanism. Nature 594, 271–276 (2021).

54. Adams, D. M. & Ricci, K. W. Vascular Anomalies: Diagnosis of Complicated Anomalies and New Medical Treatment Options. Hematol. Oncol. Clin. North Am. 33, 455–470 (2019).

55. Shaligram, S. S. et al. Bone Marrow-Derived Alk1 Mutant Endothelial Cells and Clonally Expanded Somatic Alk1 Mutant Endothelial Cells Contribute to the Development of Brain Arteriovenous Malformations in Mice. Transl. Stroke Res. (2021) doi:10.1007/s12975-021-00955-9.

56. Dumortier, J. et al. Recurrence of hereditary hemorrhagic telangiectasia after liver transplantation: clinical implications and physiopathological insights. Hepatology (2018) doi:10.1002/hep.30424.

57. George, M. A., Qureshi, S., Omene, C., Toppmeyer, D. L. & Ganesan, S. Clinical and Pharmacologic Differences of CDK4/6 Inhibitors in Breast Cancer. Front. Oncol. 11, 693104 (2021).

58. Fang, J. & Hirschi, K. Molecular regulation of arteriovenous endothelial cell specification. F1000Res. 8, (2019).

59. Chavkin, N. W. et al. Endothelial cell cycle state determines propensity for arterial-venous fate. Nat. Commun. 13, 5891 (2022).

60. Bradley, A. et al. The mammalian gene function resource: the International Knockout Mouse Consortium. Mamm. Genome 23, 580–586 (2012).

61. Sörensen, I., Adams, R. H. & Gossler, A. DLL1-mediated Notch activation regulates endothelial identity in mouse fetal arteries. Blood 113, 5680–5688 (2009).

62. Thompson, K. P. et al. Randomized, double-blind, placebo-controlled, crossover trial of oral doxycycline for epistaxis in hereditary hemorrhagic telangiectasia. Orphanet J. Rare Dis. 17, 405 (2022).

63. Elfarnawany, M. H. Signal Processing Methods for Quantitative Power Doppler Microvascular Angiography. (Western University, Electronic Thesis and Dissertation Repository. 3106. https://ir.lib.uwo.ca/etd/3106, 2015).

